# Advanced Graph Traversal used in Protein Interactions Network and Systems Biology

**DOI:** 10.1101/319624

**Authors:** Rashmi Rameshwari, Shilpa S Chapadgaonkar, T. V. Prasad

## Abstract

A methodological framework of graph traversal in Systems Biology is presented here. At present there is need to investigate system rather individual component. The proposed analysis generalizes the various idea of network representations of protein interactions. This approach highlights various methods used in construction of protein interaction graph or network using suitable algorithm. The network nodes represent protein residues. Two nodes are connected if two residues are functionally correlated during the protein interaction event. The analysis of the resulting network enables the importance of each protein for its interactions. Furthermore, the determination of the pattern of edge between residues yields insights into the function prediction of an interaction. This is of special interest to investigate intrinsically disordered proteins, since it is difficult to determine structural (three-dimensional) architecture of each proteins in protein interactions network. In present work various approaches for protein interactions network construction, models and methods along with graph theories has been discussed which can be used to reveal hidden properties and features of a network. Further effective algorithm for visualization of protein interactions is suggested. As construction of Biological network is dependent on various properties of graph. A holistic approach such as Systems Biology approach can better solve the problem. This network profiling combined with knowledge extraction will help biologist to explore hidden information in genome as well as in proteome..

## Introduction

Biological Networks are getting more advanced due to advancement in computational algorithm. Due to this more meaningful information can be extracted. These visualizations techniques are based on certain algorithm and graph theories. A graph is a perfect tool to simplify complex network properties. Graph theories are powerful tools, widely exploited in network construction and visualization of Genomics and proteomics datasets. To simplify work of a researcher different databases are categorised. It helps researcher to find out key molecule in large network.

There are various systems in nature which can be described by complex network. Starting from natural entity such as cell to physical entity such as internet, network systems are used to transmit messages from one end to another. The aim of network analysis and comparison is to identify set of entities that carry specific information to perform particular task. Topology of network varies as size of graph increases.

Since protein-protein interaction networks in which two proteins might be evolutionary related, co-occur in the literature or co-express in some experiments, resulting by this way in three different connections, each one with a different meaning (Huber et al., 2007). An example of PPI database that takes into account the different types of interactions between proteins is String (Jensen et al., 2009) and Biogrid (Stark et al., 2006). Many of these diverse network have been recently found to display network motif (Milo et al., 2002). A network motif can be referred as sub graph which occurs frequently in a network. The network of protein-protein interactions are induced by interactions between pair of high affinity sites on the protein sequences i.e. motif or domain (Ingram, Stumpf & Stark, 2006). These motif represents conserved pattern of protein sequence and are likely to interact multiple times in a protein interaction network. When a complex pattern is observed in a network it is called network motifs. It gives various information related to signal transduction and gene regulatory network (Berg & Lassig, 2004).

There are various communication process within a cell, which can be interpreted using graph network. Signal transduction is one such process within a cell used to coordinate with environmental change. This pathway is considered as directed network of reaction where stimulus binds with receptor on cell membrane. Gene regulation is controlled by generegulatory network where nodes represents genes and directed edge represent regulators. Usually these networks are represented by directed graph.

**Table 1:** Comparision of various Protein Interactions Graph traversal Algorithm.

| S.No | Features | Scholtens and Gentleman's Algorithm | Klamt and Gilles' Algorithm | Bellman–Ford's algorithm | Floyd–Warshall algorithm | Dijkstra's Algorithm |
| --- | --- | --- | --- | --- | --- | --- |
| 1 | Graph Connections | Three types of complex can be predicted:<br>a) more than one bait and more than one edge in the subgraph<br>b) single-bait-multi-hit (SBMH) complexes containing one bait and a collection of hits.<br>c) unreciprocated bait-bait (UnRBB) complexes containing two proteins, both used as baits, connected by one unreciprocated edge | Displayed using Hypergraphs which are generalized graphs, where an edge (reaction) can link $k$ nodes (reactants) with $1$ nodes (products), whereas in graphs only $1 : 1$-relations are allowed. | Computes shortest path from single source vertex to all of the other vertices in weighted digraph | It is used in finding shortest paths in a weighted graph | It is used for determination of shortest path in weighted unsigned graph |
| 2 | Algorithm Applied | Yes (Complex membership Estimation Algorithm) | Yes (Minimal cut set) | Yes (Shortest path algorithm) | Yes | Yes |
| 3 | Data set used | Publicly available AP-MS data protein Interaction | Metabolic pathway data is used. | Metabolic pathway data is used. | Genomic and proteomic dataset | Protein Interaction data |
| 4 | Applications in Biological Network | Can predict or investigate communication between cellular systems by | Used in target identification in drug discovery as well as | Used in reverse engineering of metabolic pathway | is a good choice for computing paths between all pairs of | Good for computing single source problem |
|  |  | ascertaining complex comembership for proteins known to be associated with different pathways. | for metabolic engineering towards rational strain design. |  | vertices in dense graph and large graph. |  |

In Metabolic and biochemical network, series of chemical reactions take place within a cell at different time points. These can be represented by networks which in turn is based on various graph theories. Network properties is explored using graph properties. The network can be drawn using directed, undirected, weighted and bipartite graph. As enzyme is the key component which catalyzes various biological reactions. Usually these network can be represented by bipartite graph. In undirected graph the number of connections of node to its neighbouring node can vary. If a network is directed then each node has two different degree. The indegree which is the number of incoming edges to node “i” and out degree which is the number of outgoing edges from node.

The connectivity of a network is defined as

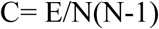

Where E= Number of Edges

N= Total number of Nodes

**FIGURE 1:**
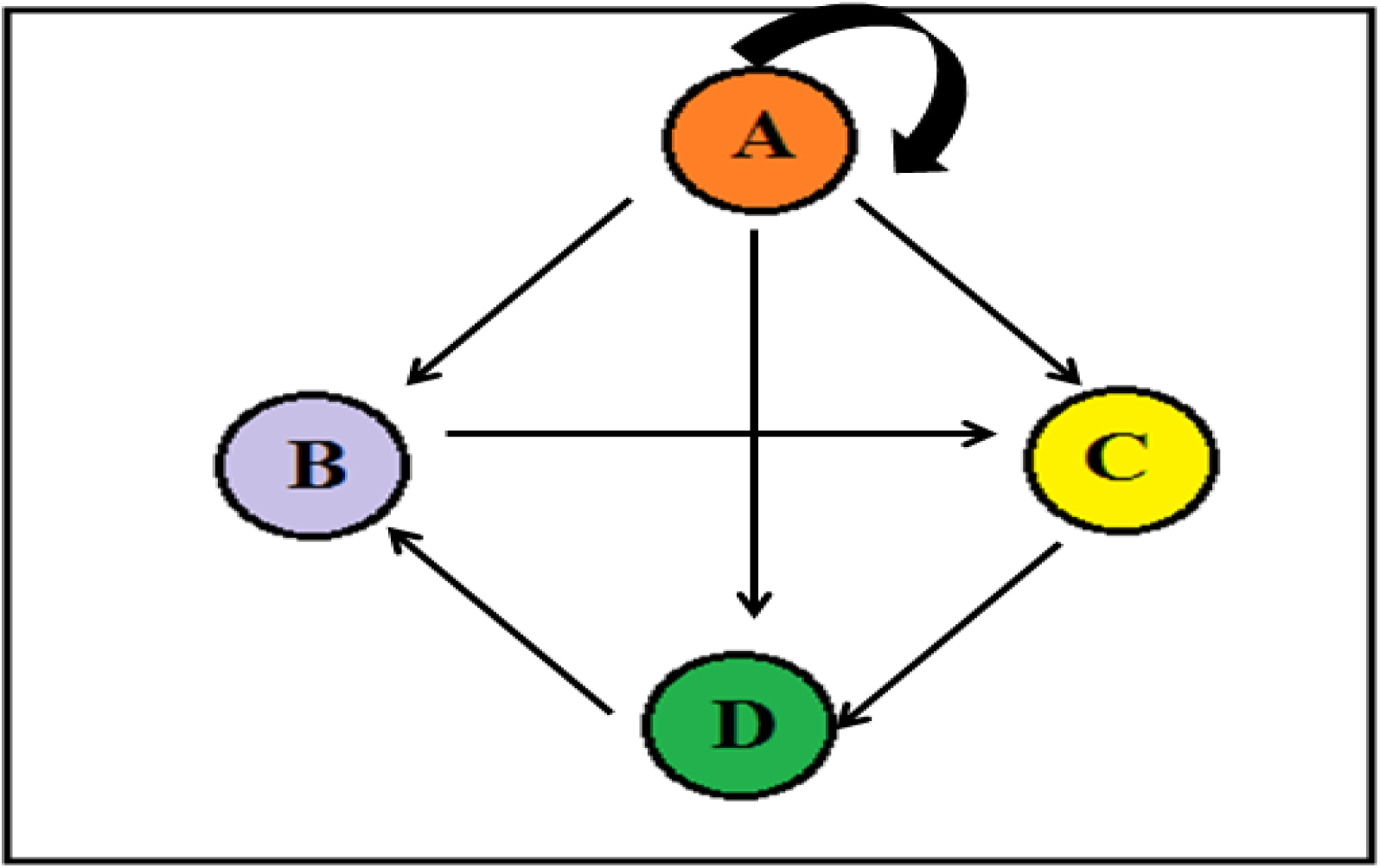
A simple graph.

Two main data structure is used to store network in graph traversal in Systems Biology. They are Adjacency matrix and Adjacency list.

Network property provides valuable information about molecule organization in a graph traversal. It gives information about evolutionary link of a molecule. It can be shown by graph density. A graph is said to be complete graph where every pair of distinct vertices is connected by a unique edge. In a graph traversal when a specific sequence of node is visited once it is called simple path and the node is not repeated in graph traversal. It can be cyclic when last node is same as first one. Sometimes in a graph traversal edge is not repeated it is called acyclic (Pavlopoulos et al., 2011) graph traversal several network model is used. For random graph Erdos-Renyi proposed a theoretical model of a network called random graph. In this graph “n” nodes joined by edges from N(N−1)/2 possible edges that have been randomly chosen (Erdos & Rényi, 1961). They gave several version of random graph.

The network with small world topology is best described by Watts and Strogatz model (Watts & Strogatz, 1998). This model describes many Biological networks like metabolic network. In this model frequency of nodes P(K) with “K” connections follows a power law distribution equation P(k) ~K ^−Y^ in which most nodes are connected with small proportions of other nodes and a small proportion of nodes are highly connected. Scale free network is best described by Barbasi-Albert mode. This model describes dynamics of almost all Biological network (Barabási & Albert, 1999).

There are many algorithms for enumeration of paths and cycle in graph traversal. However there is need of an Algorithm which can deal with incompleteness and uncertainty in graph traversal. For example, to predict a special form of protein complex data Scholtens and Gentleman (Scholtens & Gentleman, 2004) developed an algorithm. For hypergraphs, Krishnamurthy et al. describe an extension of depth first search (Krishnamurthy et al., 2003), and Klamt and Gilles developed an analog of the mincut algorithm for biochemical reaction networks (Klamt & Gilles, 2004).

For enumeration of all cycles in digraph Tarjan (Tarjan, 1972) and Johnson (Johnson, 1975) proposed algorithm. Elementary modes computation could also be used to compute path. For weighted unsigned graph Dijkstra’s algorithm is best suited (Dijkstra, 1959). For this simple breadth first search can be used. It is important for a pair of node to check whether the path is positive or negative. Although it is difficult to find whether the path is positive or negative. It is only possible in polynomial time when either the graph does not contain any negative cycles (Grotschel & Pulleyblank, 1981).

## Material and Methods

In present work graph traversal in systems biology is considered using force directed algorithm. In this hub protein (key protein for interaction) is considered as initiator of graph traversal. A graph G can be defined as a pair (V, E) where V is a set of vertices representing the nodes and E is a set of edges representing the connections between the nodes. E is defined as (i, j)| i, j Î V} the single connection between nodes i and j. In this case, it is said that i and j are neighbours. Multi-edge connection means two or more edges that have the same endpoints. Multi-edges are important for Protein interaction network construction. In this two elements can be linked by more than one connection. In such cases, each connection indicates a different type of information.

In protein interaction graph, if path starts from hub protein node which is also called seed node traverse in by breadth first or depth first. When path originates from hub protein the graph is considered as unweighted Graph while traversal in Systems Biology. In present work graph traversal is shown using force directed algorithm (‘unpublished data’, Ramehwari R et.al) in which two types of forces acts, such as attraction and repulsion forces.

According to coulomb’s Law the repulsion force acting between two object is:

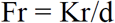

According to Hook’s Law the attraction force between them is:

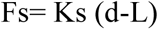

The interaction can be unidirectional when only pair of protein is involved. This mean that one protein can interact with other one protein only at a given time. But in nature this does not happen. One protein can interact with many other protein. One protein can be hub at one time while it can be hit at other time. This interactions can be visualized using various layout.

### Result

The graph traversal was studied and visualized in cancer singnaling pathway available at NetPath. Visualization was carried out by directed graph and different layout algorithm was used to show protein interaction. The different layout of graph helps in visualizing whether that hub protein is triggering other proteins or it is being triggered by any other protein. The directed graph helps in understanding key protein in protein interaction network.

**FIGURE 2:**
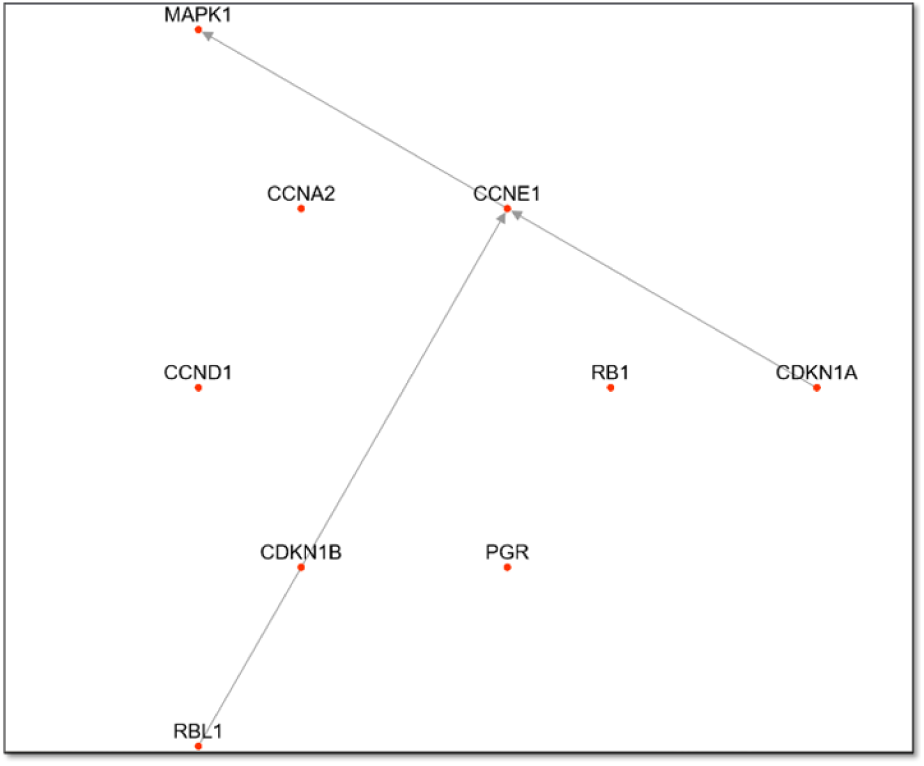
Breadth First Algorithm.

**FIGURE 3:**
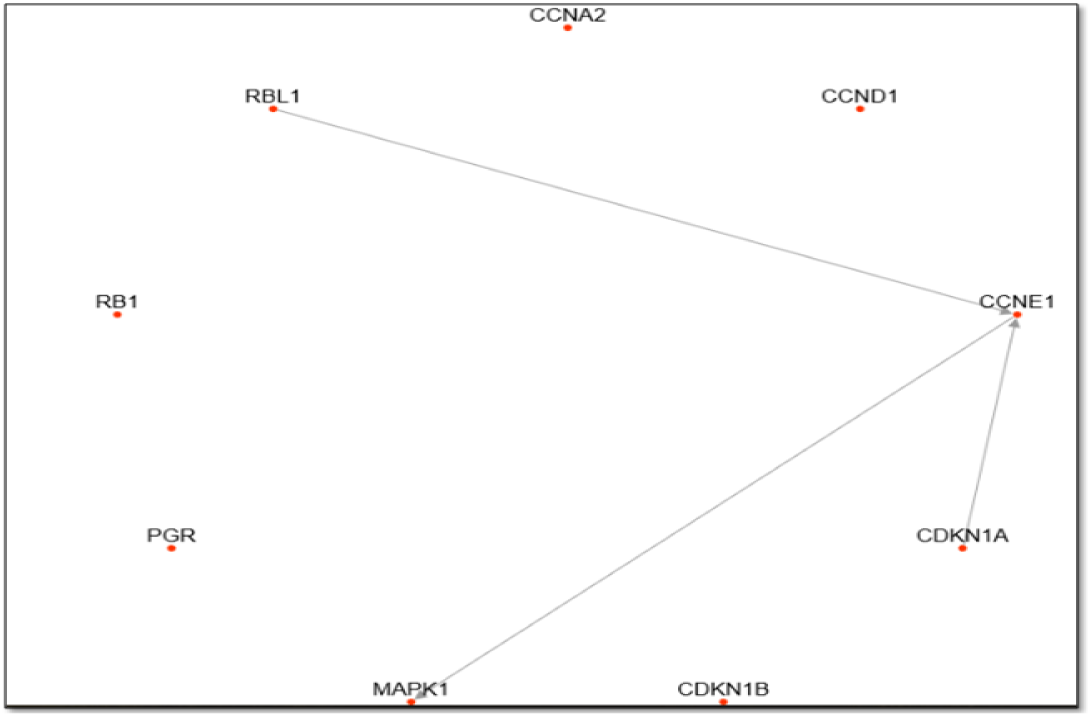
Circular Layout Algorithm.

**FIGURE 4:**
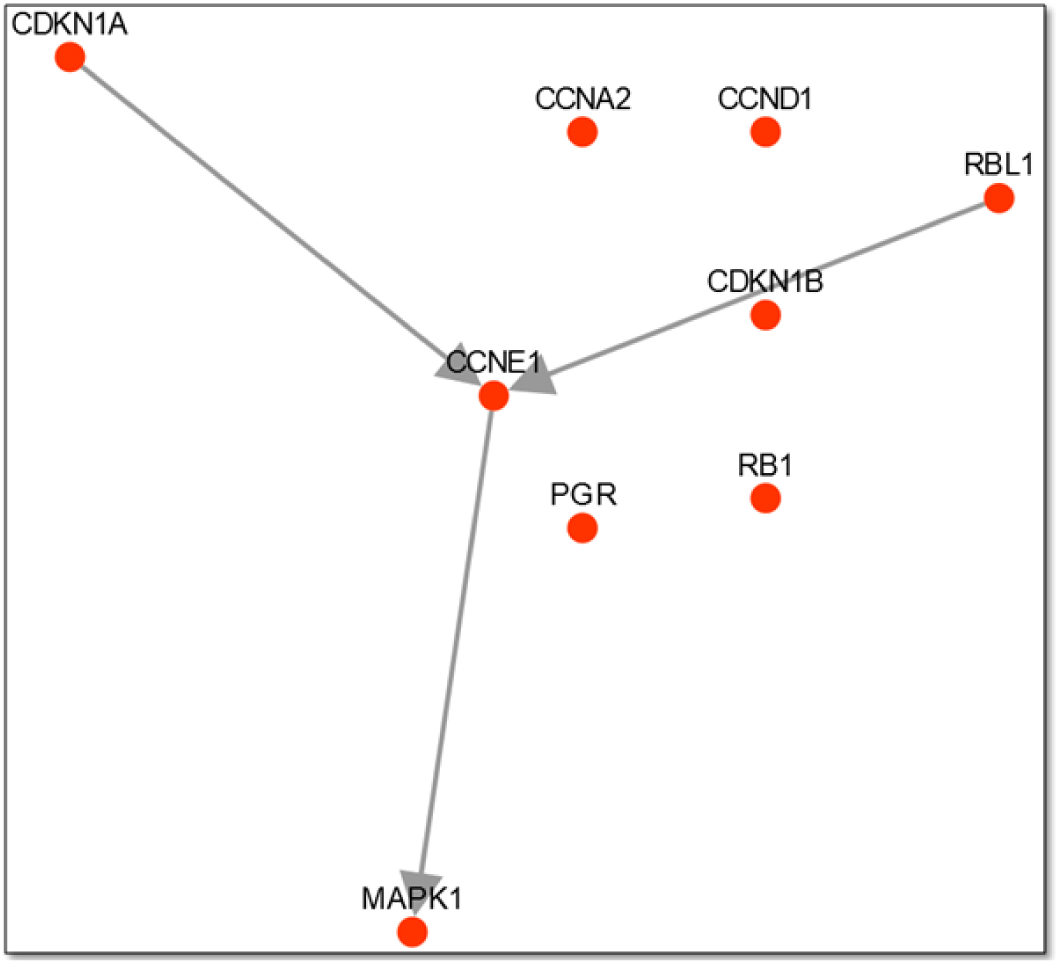
Force Directed Layout.

## Discussion

For layout of graph force directed layout algorithm was chosen. A protein physically interacts to other protein only when they share common domain. So when a protein interacts with many other proteins it is likely that it shares common domains with interacting partner. So protein localization is very important role in protein interaction. Evolutionary conservation of protein feature plays important role in protein interaction. The force which acts on a protein to interact with other is likely to be same in nature of protein. If two proteins are near by two forces act on this based on nature of protein.

## Conclusion

The listing of paths and cycles and the calculation of shortest positive or negative paths in interaction graphs are fundamental issues in Systems Biology. In the present work different algorithm for shortest path was compared. Different technique was borrowed from previous work. It is well understood that algorithms exploiting explicitly the graph structure (where each edge connects two nodes). Apart from full enumeration of paths and cycles, identification and calculation of shortest positive or negative paths and cycles in interaction graphs is a key problem. We proposed here graph traversal in which various layout algorithm is utilized for better visualization in dense graph. It is also suggested here that two step algorithm can be used for both computing exact results in smaller and medium-scale networks and approximations in large-scale networks. In biological networks that exact results can be obtained in networks with up to several hundreds nodes and interactions.

## References

Barabási A-L., Albert R. 1999. Emergence of Scaling in Random Networks. Science 286:509–512. DOI: 10.1126/science.286.5439.509.

Berg J., Lässig M. 2004. Local graph alignment and motif search in biological networks. Proceedings of the National Academy of Sciences 101:14689–14694. DOI: 10.1073/pnas.0305199101.

Dijkstra EW. 1959. A Note on Two Problems in Connexion with Graphs. Numer. Math. 1:269–271. DOI: 10.1007/BF01386390.

Erdos P., Rényi A. 1961. On the strength of connectedness of a random graph. Acta Mathematica Academiae Scientiarum Hungaricae 12:261–267. DOI: 10.1007/BF02066689.

Grötschel M., Pulleyblank WR. 1981. Weakly bipartite graphs and the Max-cut problem. Operations Research Letters 1:23–27. DOI: 10.1016/0167-6377(81)90020-1.

Huber W., Carey VJ., Long L., Falcon S., Gentleman R. 2007. Graphs in molecular biology. BMC Bioinformatics 8:S8. DOI: 10.1186/1471-2105-8-S6-S8.

Ingram PJ., Stumpf MPH., Stark J. 2006. Network motifs: structure does not determine function. BMC genomics 7:108. DOI: 10.1186/1471-2164-7-108.

Jensen LJ., Kuhn M., Stark M., Chaffron S., Creevey C., Muller J., Doerks T., Julien P., Roth A., Simonovic M., Bork P., von Mering C. 2009. STRING 8--a global view on proteins and their functional interactions in 630 organisms. Nucleic Acids Research 37:D412–416. DOI: 10.1093/nar/gkn760.

Johnson D. 1975. Finding All the Elementary Circuits of a Directed Graph. SIAM Journal on Computing 4:77–84. DOI: 10.1137/0204007.

Klamt S., Gilles ED. 2004. Minimal cut sets in biochemical reaction networks. Bioinformatics (Oxford, England) 20: 226–234.

Krishnamurthy L., Nadeau J., Ozsoyoglu G., Ozsoyoglu M., Schaeffer G., Tasan M., Xu W. 2003. Pathways database system: an integrated system for biological pathways. Bioinformatics (Oxford, England) 19: 930–937.

Milo R., Shen-Orr S., Itzkovitz S., Kashtan N., Chklovskii D., Alon U. 2002. Network motifs: simple building blocks of complex networks. Science (New York, N.Y.) 298:824–827. DOI: 10.1126/science.298.5594.824.

Pavlopoulos GA., Secrier M., Moschopoulos CN., Soldatos TG., Kossida S., Aerts J., Schneider R., Bagos PG. 2011. Using graph theory to analyze biological networks. BioData Mining 4:10. DOI: 10.1186/1756-0381-4-10.

Scholtens D., Gentleman R. 2004. Making sense of high-throughput protein-protein interaction data. Statistical Applications in Genetics and Molecular Biology 3:Article39. DOI: 10.2202/1544-6115.1107.

Stark C., Breitkreutz B-J., Reguly T., Boucher L., Breitkreutz A., Tyers M. 2006. BioGRID: a general repository for interaction datasets. Nucleic Acids Research 34:D535–539. DOI: 10.1093/nar/gkj109.

Tarjan RE. 1972. Enumeration of the Elementary Circuits of a Directed Graph.

Watts DJ., Strogatz SH. 1998. Collective dynamics of “small-world” networks. Nature 393:440–442. DOI: 10.1038/30918.

